# Rapid computations of spectrotemporal prediction error support perception of degraded speech

**DOI:** 10.1101/2020.04.22.054726

**Authors:** Ediz Sohoglu, Matthew H. Davis

## Abstract

Human speech perception can be described as Bayesian perceptual inference but how are these Bayesian computations instantiated neurally? We use magnetoencephalographic recordings of brain responses to degraded spoken words as a function of signal quality and prior knowledge to demonstrate that spectrotemporal modulations in speech are more clearly represented in neural responses than alternative speech representations (e.g. spectrogram or articulatory features). We found an interaction between speech signal quality and expectations from prior written text on the quality of neural representations; increased signal quality enhanced neural representations of speech that mismatched with prior expectations, but led to greater suppression of speech that matched prior expectations. This interaction is a unique neural signature of prediction error computations and already apparent in neural responses within 250 ms of speech input. Our findings contribute towards the detailed specification of a computational model of speech perception based on predictive coding frameworks.

## Introduction

Although we understand spoken language rapidly and automatically, speech is an inherently ambiguous acoustic signal, compatible with multiple interpretations. Such ambiguities are evident even for clearly spoken speech: A /t/ consonant will sometimes be confused with /p/, as these are both unvoiced stops with similar acoustic characteristics (Warner et al., 2014). In real-word environments, where the acoustic signal is degraded or heard in the presence of noise or competing speakers, additional uncertainty arises and speech comprehension is further challenged (Mattys et al., 2012; Peelle, 2018).

Given the uncertainty of the speech signal, listeners must exploit prior knowledge or expectations to constrain perception. For example, ambiguities in perceiving individual speech sounds are more readily resolved if those sounds are heard in the context of a word (Ganong, 1980; Rogers and Davis, 2017). Following probability theory, the optimal strategy for combining prior knowledge with sensory signals is by applying Bayes theorem to compute the posterior probabilities of different interpretations of the input. Indeed, it has been suggested that spoken word recognition is fundamentally a process of Bayesian inference (Norris and McQueen, 2008). The present work aims to establish how these Bayesian computations might be instantiated neurally.

There are at least two representational schemes by which the brain could implement Bayesian inference (depicted in Figure 1C; see Aitchison and Lengyel, 2017). One possibility is that neural representations of sensory signals are enhanced or ‘sharpened’ by prior knowledge (Murray et al., 2004; Friston, 2005; Blank and Davis, 2016; de Lange et al., 2018). Under a sharpening scheme, neural responses directly encode posterior probabilities and hence representations of speech sounds are enhanced in the same way as perceptual outcomes are enhanced by prior knowledge (McClelland and Elman, 1986; McClelland et al., 2014). Alternatively, neural representations of expected speech sounds are subtracted from bottom-up signals, such that only the unexpected parts (i.e. ‘prediction error’) are passed up the cortical hierarchy to update higher-level representations (Rao and Ballard, 1999). Under this latter representational scheme, higher-level neural representations come to encode posterior probabilities (as required by Bayesian inference) but this is achieved by an intermediate process in which prediction errors are computed (Aitchison and Lengyel, 2017). In many models, both representational schemes are utilized, in separate neural populations (cf. predictive coding, Rao and Ballard, 1999; Spratling, 2008; Bastos et al., 2012) or at different times after stimulus onset (Press et al., 2020).

**Figure 1.**
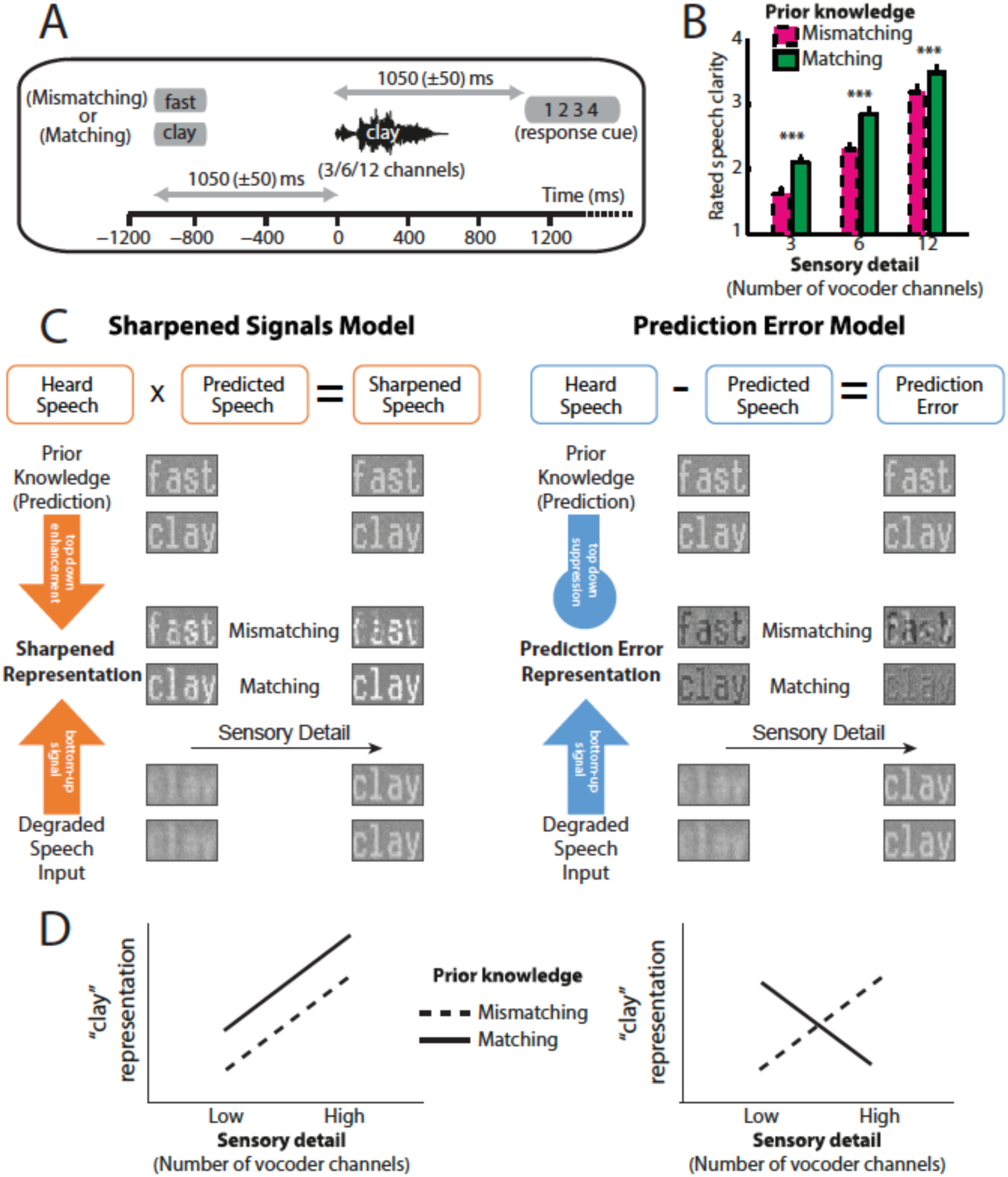
Overview of experimental design and hypotheses. A) On each trial, listeners heard and judged the clarity of a degraded spoken word. Listeners’ prior knowledge of speech content was manipulated by presenting matching (“clay”) or mismatching (“fast”) text before spoken words presented with varying levels of sensory detail (3/6/12-channel vocoded). B) Ratings of speech clarity were enhanced not only by increasing sensory detail but also by prior knowledge from matching text (graph reproduced from Sohoglu and Davis, 2016). C) Schematic illustrations of two representational schemes by which prior knowledge and speech input are combined. For illustrative purposes, we depict degraded speech as visually degraded text. Under a sharpening scheme (left panels), neural representations of degraded sensory signals (bottom) are enhanced by matching prior knowledge (top) in the same way as perceptual outcomes are enhanced by prior knowledge. Under a prediction error scheme (right panels), neural representations of expected speech sounds are subtracted from sensory signals. These two accounts make different predictions for experiments that assess the content of neural representations when sensory detail and prior knowledge of speech are manipulated. D) Theoretical predictions for sharpened signal (left) and prediction error (right) models. In a sharpened signal model, representations of the heard spoken word (e.g. “clay”) are most accurately encoded in neural responses when increasing speech sensory detail and matching prior knowledge combine to enhance perception. Conversely, for models that represent prediction error, an interaction between sensory detail and prior knowledge is observed. For speech that mismatches with prior knowledge, increasing sensory detail results in better encoding of the heard word “clay” because bottom-up input remains unexplained. Conversely, for speech that matches prior knowledge, increased sensory detail results in worse encoding of “clay” because bottom-up input is explained away.

A range of experimental evidence has been used to distinguish sharpened and prediction error representations. An often observed finding is that matching prior knowledge reduces the amplitude of evoked neural responses (Ulanovsky et al., 2003; Grill-Spector et al., 2006; Rabovsky et al., 2018). This observation is commonly attributed to prediction errors since expected stimuli are relatively predictable and therefore should evoke reduced prediction error. However, reduced activity is equally compatible with sharpened responses because under this representational scheme, neuronal activity encoding competing features (i.e. ‘noise’) is suppressed (see Figure 1C; Murray et al., 2004; Friston, 2005; Blank and Davis, 2016; Aitchison and Lengyel, 2017; de Lange et al., 2018).

One way to adjudicate between representations is by manipulating signal quality alongside prior knowledge and measuring the consequences for the pattern (rather than only the mean) of neural responses. Computational simulations reported by Blank and Davis (2016) demonstrate a unique hallmark of prediction errors which is that neural representations of sensory stimuli show an interaction between signal quality and prior knowledge (see Figure 1D). This interaction arises because sensory signals that match prior expectations are explained away more effectively as signal quality increases and hence neural representations are suppressed even as perceptual outcomes improve. Whereas for sensory signals that follow uninformative prior expectations, increased signal quality leads to a corresponding increase in sensory information that remains unexplained (see Figure 1C). In contrast, in computational simulations implementing a sharpening scheme, representational patterns are similarly enhanced by increased signal quality and matching prior knowledge (see Figure 1D; Murray et al., 2004; Friston, 2005; Blank and Davis, 2016; Aitchison and Lengyel, 2017; de Lange et al., 2018). Using prior written text to manipulate listeners’ prior knowledge of degraded spoken words, Blank and Davis (2016) showed that multivoxel representations of speech in the superior temporal gyrus (as measured by fMRI) showed an interaction between signal quality and prior expectations, consistent with prediction error computations. This is despite the observation that mean multivoxel responses to speech were always reduced by matching expectations, no matter the level of sensory detail.

Although the study of Blank and Davis (2016) provides evidence in support of prediction errors, key questions remain. Firstly, it remains unclear at which levels of representation prediction errors are computed. Predictive coding models that utilize prediction errors are hierarchically organized such that predictions are signaled by top-down connections and prediction errors by bottom-up connections. Therefore, in principle, prediction errors will be computed at multiple levels of representation. Previous studies using similar paradigms have suggested either a higher-level phonetic (Di Liberto et al., 2018a) or a lower-level acoustic locus of prediction error representations (Holdgraf et al., 2016). Secondly, due to the sluggishness of the BOLD signal, the timecourse over which prediction errors are computed is unknown. Therefore, it is unclear whether prediction errors are formed only at late latencies following re-entrant feedback (> 250 ms) or more rapidly during the initial feedforward sweep of cortical processing (Sohoglu and Davis, 2016; Kok et al., 2017; de Lange et al., 2018; Di Liberto et al., 2018a).

In the current study, we reanalyzed MEG recordings of neural activity from a previous experiment in which we simultaneously manipulated signal quality and listeners’ prior knowledge during speech perception (Sohoglu and Davis, 2016). Listeners heard degraded spoken words with varying amounts of sensory detail and hence at different levels of signal quality. Before each spoken word, listeners’ read matching or mismatching text and therefore had accurate or inaccurate prior knowledge of upcoming speech content (Figure 1A). Our previously reported analyses focused on the mean amplitude of evoked responses, which as explained above, cannot adjudicate between sharpened and prediction error representations. We therefore used linear regression to test which of several candidate speech features are encoded in MEG responses (Ding and Simon, 2012; Pasley et al., 2012; Crosse et al., 2016; Holdgraf et al., 2017) and further asked how those feature representations are modulated by signal quality and prior knowledge. Following Blank and Davis (2016), a two-way interaction between sensory detail and prior knowledge is diagnostic of prediction errors (see Figure 1C and D). Because of the temporal resolution of MEG, if we observe such an interaction we can also determine whether prediction errors are computed at an early latency of processing (< 250 ms).

## Results

### Behaviour

During the MEG recordings, listeners completed a clarity rating task in which they judged the subjective clarity of each degraded spoken word (Figure 1A and 1B). Ratings of speech clarity were enhanced both when sensory detail increased (*F* (2,40) = 295, *p* < .001) and when listeners had prior knowledge from matching written text (*F* (1,20) = 93.2, *p* < .001). These behavioural results have previously been reported (Sohoglu and Davis, 2016) but we include them here to facilitate interpretation of the present MEG analyses.

### Encoding analysis: Stimulus feature space selection

We used ridge regression to predict the MEG data from the stimulus features. A model that accurately predicts the MEG data would indicate that the component features in the model are well represented in neural responses. Four feature spaces were obtained from the original clear versions of the spoken stimuli (i.e. prior to noise-vocoding; see Figure 2). In the first stage of our analysis, we examined which of these four feature spaces best predicted the MEG data.

**Figure 2.**
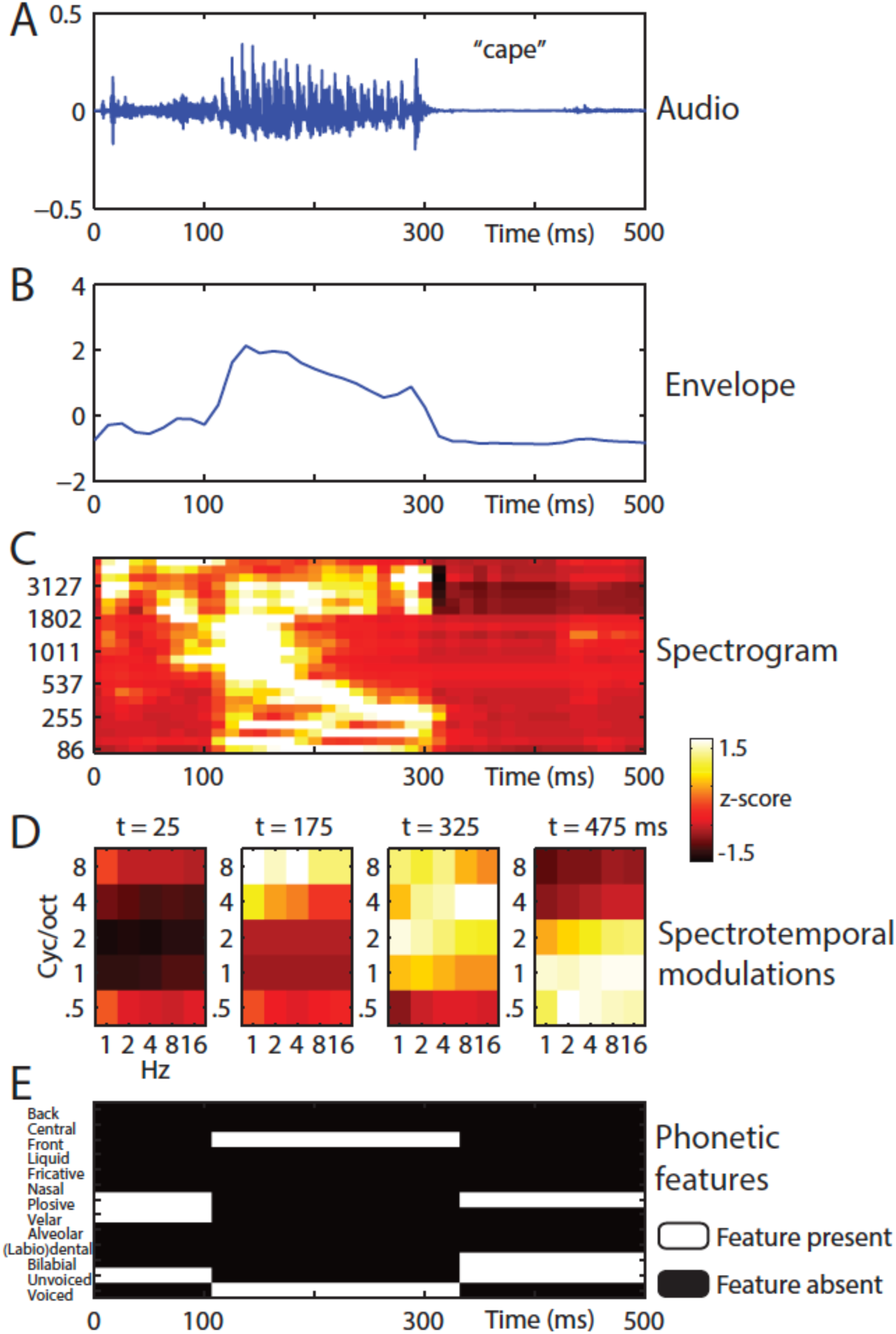
Stimulus feature spaces used to model MEG responses, for the example word “cape”, A) shown as an audio waveform for the original clear recording (i.e. prior to vocoding). B) Envelope: broadband envelope derived from summing the envelopes across all spectral channels of a 24-channel noise-vocoder. C) Spectrogram: derived from the envelope in each spectral channel of a 24-channel noise-vocoder. D) Spectrotemporal modulations: derived from the spectral and temporal decomposition of a spectrogram into 25 spectrotemporal channels, illustrated at regular temporal intervals through the spoken word. E) Phonetic features: derived from segment-based representations of spoken words.

We averaged model accuracies over conditions and over the 20 sensors with the highest model accuracies (computed separately for each feature space, hemisphere and participant; shown in Figure 3A). There was a significant main effect of feature space on model accuracies (*F* (3,60) = 51.5, *p* < .001), which did not interact with hemisphere (*F* (3,60) = 1.56, *p* = .209). Post-hoc *t*-tests revealed that the spectrotemporal modulation feature space best predicted the MEG data (*p*-values shown in Figure 3A). Sensor selections (i.e. those sensors with the highest model accuracies; shown in Figure 3B) were similarly distributed over temporal and frontal sites for all feature spaces, consistent with bilateral neural generators in superior temporal cortex.

**Figure 3.**
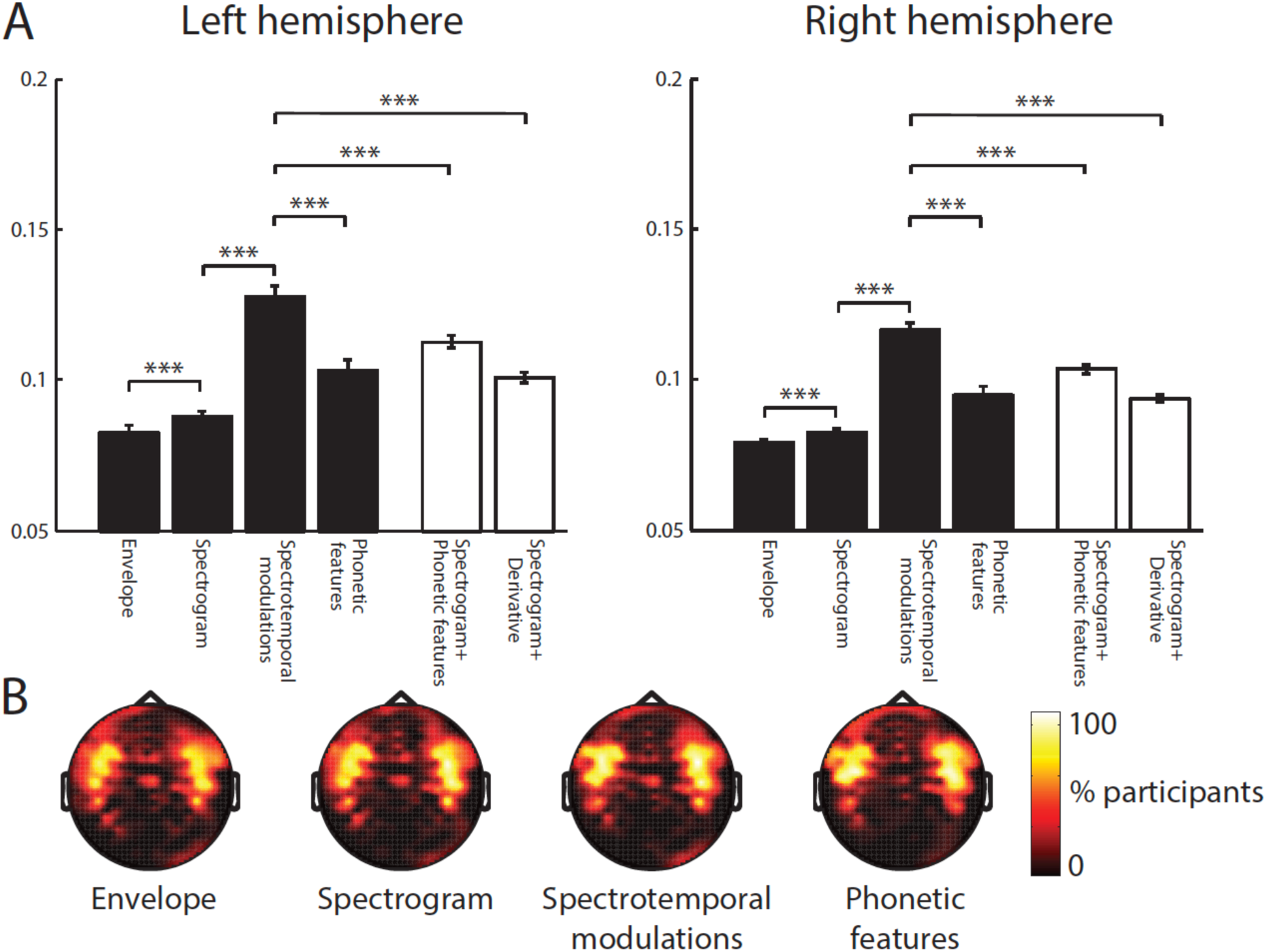
Encoding model comparison. A) Mean model accuracies for the four feature spaces (black bars) and the two feature space combinations (white bars). Error bars represent standard error of the mean after removing between-subject variance, suitable for repeated-measures comparisons (Loftus and Masson, 1994). Braces indicate significance of paired t-tests *** p < .001 B) Topographic distribution of MEG gradiometer sensors over which model accuracies were averaged. In each hemisphere of each participant, we selected 20 sensors with the highest model accuracies. The topographies show the percentage of participants for which each sensor was selected, consistent with neural sources in superior temporal cortex.

We additional tested more complex combinations of feature spaces that previous work (Di Liberto et al., 2015; Daube et al., 2019) has shown to be good models of neural responses (shown in Figure 3A). The first of these models combined the spectrogram with phonetic features while the second model combined the spectrogram with the half-wave derivative of the spectrogram. We replicated previous findings in that both the Spectrogram+Phonetic features model (*F* (1,20) = 52.8, *p* < .001) and the Spectrogram+Derivative model (*F* (1,20) = 110, *p* < .001) outperformed the spectrogram feature space. However, the spectrotemporal modulation feature space remained the model that best predicted neural responses, outperforming Spectrogram+Phonetic features (*F* (1,20) = 16.5, *p* < .01) and Spectrogram+Derivative (*F* (1,20) = 98.2, *p* < .001). For completeness, we also report that Spectrogram+Phonetic features outperformed Spectrogram+Derivative (*F* (1,20) = 16.1, *p* < .01). No effects involving hemisphere were significant (all *p*’s > .125).

### Acoustic analysis: Stimulus modulation content and effect of vocoding

Having identified spectrotemporal modulations as the feature space that is most accurately represented in neural responses, we next characterized the spectrotemporal modulations that convey speech content and how those modulations are affected by noise-vocoding with a wide range of spectral channels, from 1 to 24 channels. As shown in Figure 4A, modulations showed a lowpass profile and were strongest in magnitude for low spectrotemporal frequencies. This is consistent with previous work (Voss and Clarke, 1975; Singh and Theunissen, 2003) demonstrating that modulation power of natural sounds decays with increasing frequency following a 1/f relationship (where f is the frequency). Within this lowpass region, different spectrotemporal modulations relate to distinct types of speech sound (Elliott and Theunissen, 2009). For example, fast temporal and broad spectral modulations reflect transient sounds such as stop consonant release bursts whereas slow temporal and narrowband spectral modulations reflect sounds with a sustained spectral structure such as vowels. More intermediate modulations correspond to formant transitions that cue consonant place and manner of articulation (Liberman et al., 1967).

**Figure 4.**
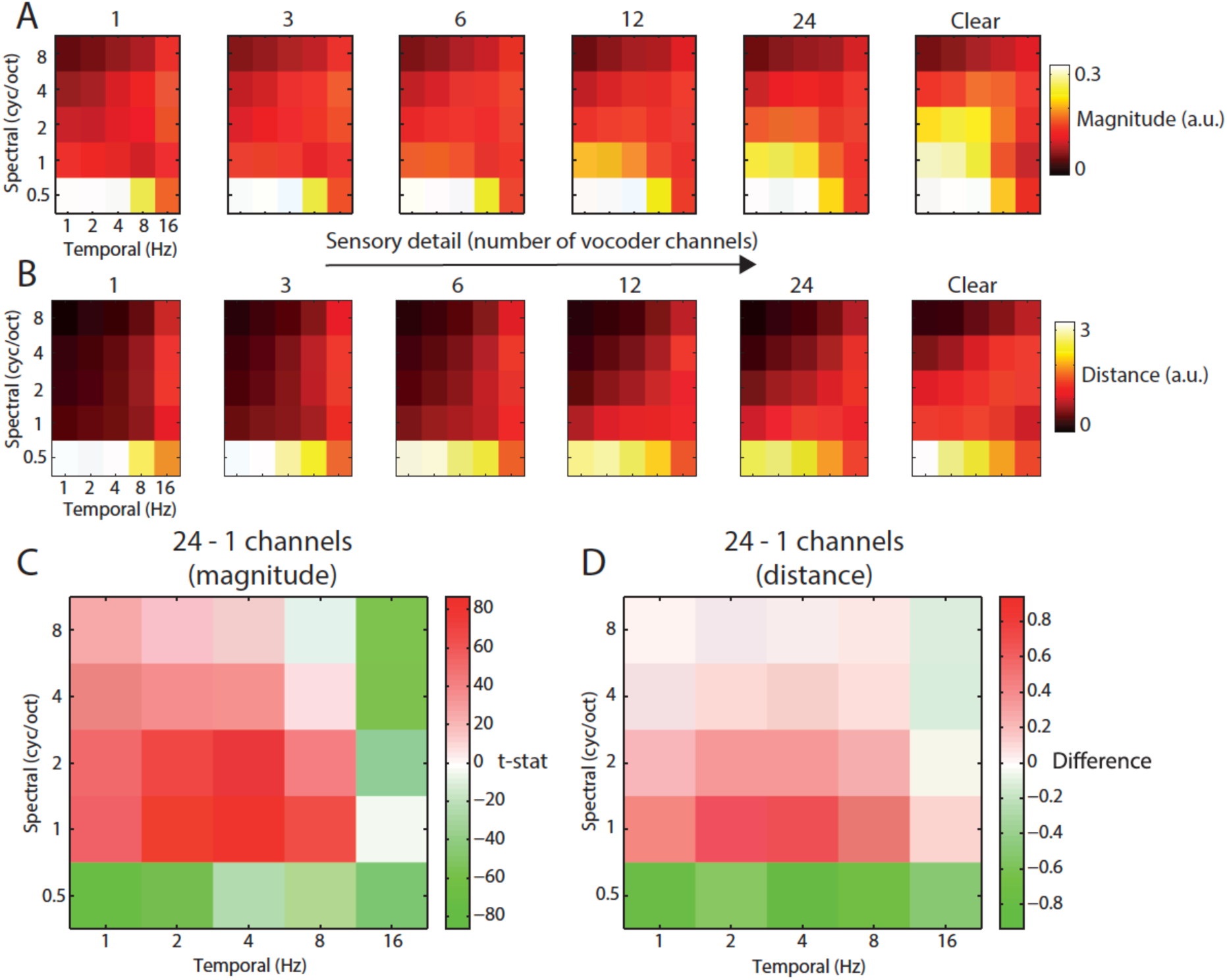
Acoustic comparison of vocoded speech stimuli at varying levels of sensory detail. A) The magnitude of different spectrotemporal modulations for speech vocoded with different numbers of channels and clear speech (for comparison only; clear speech was not presented to listeners in the experiment). B) Mean between-word Euclidean distance for different spectrotemporal modulations in speech vocoded with different numbers of channels. C) Paired t-test showing significant differences in spectrotemporal modulation magnitude for comparison of 958 spoken words vocoded with 24 channels versus 1 channel (p < .05 FDR corrected for multiple comparisons across spectrotemporal modulations). D) Mean difference of between-word Euclidean distances for different spectrotemporal modulations for 24 versus 1 channel vocoded speech.

Over items, increasing the number of vocoder channels resulted in significantly higher signal magnitude specific to intermediate spectral and temporal modulations (1-2 cycles per octave and 2-4 Hz, all effects shown are FDR corrected for multiple comparisons across spectrotemporal modulations; see Figure 4C). Thus, while low frequency modulations dominate the speech signal overall (irrespective of the number of vocoder channels), it is the intermediate spectrotemporal modulations that are most strongly affected by noise-vocoding. These intermediate spectrotemporal modulations are known to support speech intelligibility (Elliott and Theunissen, 2009; Venezia et al., 2016; Flinker et al., 2019), consistent with the strong impact of the number of vocoder channels on word report accuracy (e.g. Shannon et al., 1995; Davis and Johnsrude, 2003; Scott et al., 2006; Obleser et al., 2008) and subjective clarity (e.g. Obleser et al., 2008; Sohoglu et al., 2014). The opposite effect (i.e. *decreased* signal magnitude with an increasing number of vocoder channels) was observed for broadband spectral modulations (0.5 cycles/octave) across all temporal modulation rates (reflecting the increase in envelope co-modulation when fewer spectral channels are available) and for fast temporal modulations (16 Hz) and narrowband (> 2 cycles/octave) spectral modulations (reflecting stochastic fluctuations in the noise carrier; Stone et al., 2008).

We also asked which spectrotemporal modulations were most informative for discriminating between different words. For each spectrotemporal modulation bin, we computed the Euclidean distances between the timeseries of all the words in our stimulus set and averaged the resulting distances (see Figure 4B). Between-word distances resembled the magnitude-based analysis, with greatest distance for low spectrotemporal frequencies. Between-word distances also showed a similar effect of increasing the number of vocoder channels (see Figure 4D).

### Encoding analysis: Condition effects

We next determined between-condition differences in the ability of the spectrotemporal modulation feature space to predict MEG responses. Any such differences would indicate that the neural representation of spectrotemporal modulations is modulated by prior knowledge or speech sensory detail. Model accuracies for different conditions are shown in Figure 5A. There was a significant interaction between prior knowledge and speech sensory detail (*F* (2,40) = 5.41, *p* = .01), which marginally interacted with hemisphere (*F* (2,40) = 3.57, *p* = .051). Follow-up tests in left hemisphere sensors again showed a statistical interaction between sensory detail and prior knowledge (*F* (2,40) = 7.66, *p* < .01). For speech that Mismatched with prior knowledge, model accuracies increased with increasing speech sensory detail (*F* (2,40) = 4.49, *p* < .025). In the Matching condition, however, model accuracies decreased with increasing sensory detail (*F* (2,40) = 3.70, *p* < .05). The statistical interaction between sensory detail and prior knowledge can also be characterized by comparing model accuracies for Matching versus Mismatching conditions at each level of sensory detail. While model accuracies were greater for Matching versus Mismatching conditions for low clarity 3 channel speech (*t* (20) = 2.201, *p* < .05), the opposite was true for high clarity 12 channel speech (*t* (20) = 2.384, *p* < .05). Model accuracies did not significantly differ between Matching and Mismatching conditions for 6 channel speech with intermediate clarity (*t* (20) = 1.085, *p* = .291). This interaction is consistent with the prediction error account and inconsistent with sharpened representations (compare Figure 5A with Figure 1D). No significant differences were observed in the right hemisphere.

**Figure 5.**
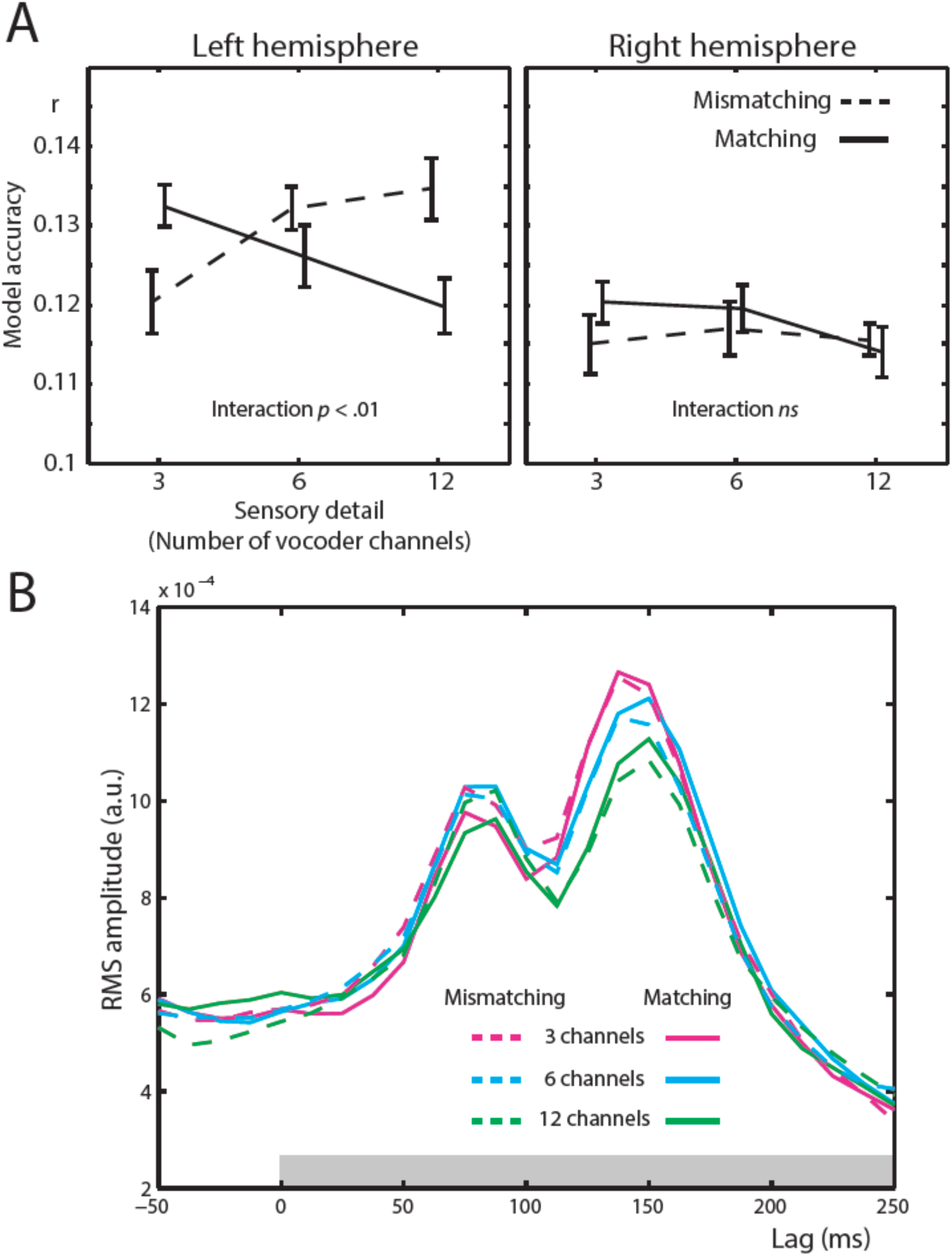
Spectrotemporal modulation encoding model results. A) Mean model accuracies for lags between 0 and 250 ms as a function of sensory detail (3/6/12 channel vocoded words), and prior knowledge (speech after mismatching/matching text) for left and right hemisphere sensors. B) Root Mean Square (RMS) amplitude across all left hemisphere sensors for the Temporal Response Functions (TRFs) for each condition, averaged over spectrotemporal modulations and participants. The gray box at the bottom of the graph indicates the lags used in computing the model accuracy data in panel A.

Examination of the weights of the spectrotemporal modulation model i.e. temporal response functions (TRFs) show two peaks at 87.5 and 150 ms (shown in Figure 5B). This indicates that variations in spectrotemporal modulations are linked to the largest changes in MEG responses 87.5 and 150 ms later. These findings are consistent with previous work showing multiple early components in response to ongoing acoustic features in speech that resemble the prominent P1, N1 and P2 components seen when timelocking to speech onset (e.g. Lalor and Foxe, 2010; Ding and Simon, 2012; Di Liberto et al., 2015). Thus, analysis of the TRFs confirms that encoding of spectrotemporal modulations is associated with short-latency neural responses at a relatively early hierarchical stage of speech processing.

### Decoding analysis

To link MEG responses with specific spectrotemporal modulations, we used linear regression for data prediction in the opposite direction: from MEG responses to speech spectrotemporal modulations (i.e. decoding analysis). As shown in Figure 6A, intermediate temporal modulations (2-4 Hz) were best decoded from MEG responses. This observation is consistent with previous neurophysiological (Ahissar et al., 2001; Luo and Poeppel, 2007; Peelle et al., 2012; Ding and Simon, 2014; Di Liberto et al., 2015; Park et al., 2015; Obleser and Kayser, 2019) and fMRI (Santoro et al., 2017) data showing that intermediate temporal modulations are well-represented in auditory cortex. These intermediate temporal modulations are also most impacted by noise-vocoding (see Figure 4C and D) and support speech intelligibility (Elliott and Theunissen, 2009; Venezia et al., 2016).

**Figure 6.**
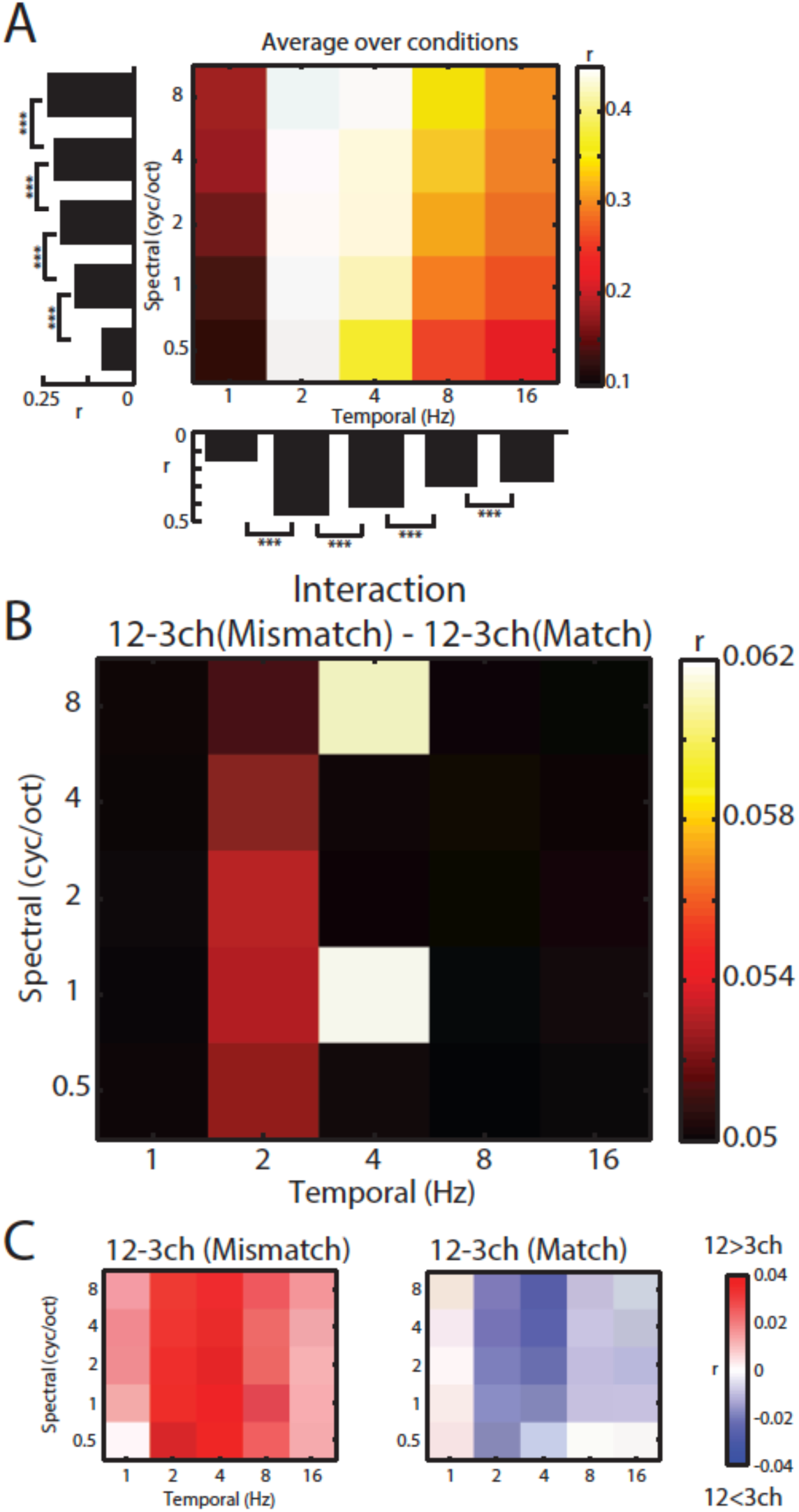
Decoding of spectrotemporal modulations from MEG responses to speech. A) The grid shows model accuracies for specific spectrotemporal modulations averaged over conditions. Left bar graph depicts model accuracy for each spectral modulation frequency, averaged over temporal modulations. Bottom bar graph depicts model accuracy for each temporal modulation frequency, averaged over spectral modulations. Braces indicate significance of paired t-tests *** p < .001 B) Effect size (model accuracy differences, r) for the interaction contrast: 12-3 channels (Mismatch) – 12-3 channels (Match). The effect size display has been thresholded so as to only show cells in which the sensory detail by prior knowledge interaction is statistically significant at *p* < .05 FDR corrected for multiple comparisons across spectrotemporal modulations. C) Effect size (model accuracy differences, r) for comparisons between 12 and 3 channels, computed separately for Mismatching and Matching conditions. Red shows greater model accuracy for 12 channel than for 3 channel speech (observed for speech that Mismatches with written text). Blue shows lower model accuracy for 12 channel than for 3 channel speech (observed for speech that matches written text). ch = channels

Having previously identified a significant interaction between prior knowledge and sensory detail in our encoding analysis, we next conducted a targeted *t*-test on decoding accuracies using the interaction contrast: 12-3 channels (Mismatching) – 12-3 channels (Matching). This allowed us to identify *which* spectrotemporal modulations show evidence of the interaction between prior knowledge and sensory detail. As shown in Figure 6B, this interaction was observed at intermediate (2-4 Hz) temporal modulations (FDR corrected across spectrotemporal modulations). Visualization of the difference between 12 and 3 channels for Mismatching and Matching conditions separately confirmed that this interaction was of the same form observed previously (i.e. increasing sensory detail led to opposite effects on decoding accuracy when prior knowledge mismatched versus matched with speech). Thus, intermediate temporal modulations are well represented in MEG responses and it is these representations that are affected by our manipulations of sensory and prior knowledge.

## Discussion

Here, using linear regression and MEG responses to noise-vocoded words, we report four main findings. Firstly, spectrotemporal modulation content in the acoustic signal is well represented in MEG responses, more so than alternative speech features (envelope, spectrogram and phonetic features). Second, the information content of speech is well characterized by spectrotemporal modulations. Modulations at intermediate temporal (2-4Hz) and spectral scales (1-2 cycles/octave) are critical for distinguishing between individual spoken words and these spectrotemporal modulations are affected by noise-vocoding manipulations that impact speech intelligibility. Thirdly, signal quality and prior knowledge have an interactive influence on these modulation-based representations: when mismatching text precedes degraded speech, neural representations of spectrotemporal modulations are enhanced whereas the opposite effect is observed when text matches speech. Fourthly, this interaction is observed specifically for neural representations of intermediate spectrotemporal modulations that convey speech content. This result stands in marked contrast to what would be expected on the basis of a sharpening account in which both signal quality and prior knowledge enhance neural representations, just as they enhance perceptual clarity. An interactive influence of signal quality and prior knowledge is, however, fully consistent with prediction error representations in which neurons signal the difference between expected and heard speech (Blank and Davis, 2016).

### Cortical responses encode spectrotemporal modulations in speech

Our finding that MEG responses are driven by spectrotemporal modulations in speech adds to a growing body of evidence implicating spectrotemporal representations as an important component of the neural code by which the cortex represents speech and other sounds. Linguistically relevant features such as formant transitions and speaker fundamental frequency are clearly apparent in these modulations and their removal by filtering has marked consequences for speech intelligibility and speaker discrimination (Elliott and Theunissen, 2009; Flinker et al., 2019). In animal electrophysiology (Chi et al., 2005; Theunissen and Elie, 2014), neurons in auditory cortex are shown to be well-tuned to spectrotemporal features in the modulation domain. These findings complement human intracranial (Pasley et al., 2012; Hullett et al., 2016) and fMRI (Santoro et al., 2014, 2017) studies demonstrating that the relationship between auditory stimuli and neural responses is best modelled using spectrotemporal modulations.

Analysis of the acoustic properties of noise-vocoded spoken words show that although low-frequency spectral and temporal modulations dominate the speech signal, only the intermediate modulations (2-4 Hz temporally; 1-2 cycles per octave spectrally) are degraded by noise-vocoding that severely impairs intelligibility (e.g. Shannon et al., 1995; Davis and Johnsrude, 2003; Scott et al., 2006; Obleser et al., 2008). These intermediate spectrotemporal modulations encode formant transitions that are an important cue to consonant place and manner of articulation (Liberman et al., 1967; Roberts et al., 2011). Notably, our decoding analysis shows that it was the intermediate temporal modulations (2-4 Hz) that were preferentially represented in MEG responses. Thus, rather than faithfully encoding the acoustic structure of speech by tracking low frequency modulations, cortical responses are selectively tuned to acoustic properties (intermediate spectrotemporal modulations) that convey speech information.

Our findings agree with recent work (Daube et al., 2019) suggesting that cortical responses to speech (as measured with MEG) are not as invariant to acoustic detail as would be expected for a higher-level categorical representation such as phonetic features or phonemes (Di Liberto et al., 2015). Despite the lack of full invariance, spectrotemporal modulations are nonetheless a higher-level representation of the acoustic signal than, for example, the spectrogram. Whereas a spectrogram-based representation would faithfully encode any pattern of spectrotemporal stimulation, a modulation-based representation would be selectively tuned to patterns that change at specific spectral scales or temporal rates. Such selectivity is a hallmark of neurons in higher-levels of the auditory system (Rauschecker and Scott, 2009). Our findings are thus consistent with the notion that superior temporal neurons, which are a major source of speech-evoked MEG responses (e.g. Bonte et al., 2006; Sohoglu et al., 2012; Brodbeck et al., 2018), lie at an intermediate stage of the speech processing hierarchy (e.g. Mesgarani et al., 2014; Evans and Davis, 2015; Yi et al., 2019). It should be noted however that in Daube et al. (2019), the best acoustic predictor of MEG responses was based on spectral onsets rather than spectrotemporal modulations. It may be that the use of single words in the current study reduced the importance of spectral onsets, which may be more salient in the syllabic transitions during connected speech as used by Daube et al. (2019). However, there are many other differences in the acoustic, phonetic and lexical properties of single spoken words and connected speech – including differences in speech rate, co-articulation, segmentation and predictability. Future work is needed to examine whether and how these differences are reflected in neural representations.

### Prediction error in speech representations

Going beyond other studies, however, our work further shows that neural representations of speech sounds reflect a combination of the acoustic properties of speech, and prior knowledge or expectations. Our experimental approach – combining manipulations of sensory detail and matching or mismatching written text cues – includes specific conditions from previous behavioural (Sohoglu et al., 2014) and neuroimaging (Sohoglu et al., 2012; Blank and Davis, 2016; Sohoglu and Davis, 2016; Blank et al., 2018) studies that were designed to distinguish different computational theories of speech perception. In particular, neural representations that show an interaction between sensory detail and prior knowledge provide unambiguous support for prediction error over sharpening coding schemes (Blank and Davis, 2016). This interaction has previously been demonstrated for multivoxel patterns as measured by fMRI (Blank and Davis, 2016). The current study goes beyond this previous fMRI work in establishing the latency at which prediction errors are computed. As prediction errors reflect the outcome of a comparison between top-down and bottom-up signals, it has been unclear whether such a computation occurs only at late latencies (>250 ms after the speech signal) and plausibly results from re-entrant feedback connections that modulate late neural responses (Sohoglu and Davis, 2016; Kok et al., 2017; de Lange et al., 2018; Di Liberto et al., 2018a). However, because the fit of our regression models to MEG data was assessed using lags from 0 to 250 ms relative to the acoustic signal, the neural interaction between sensory and prior knowledge that we observe must emerge within this latency range. This suggests that prediction errors are computed during early feedforward sweeps of processing and are tightly linked to ongoing changes in speech input.

Previous electrophysiological work (Holdgraf et al., 2016; Di Liberto et al., 2018b) has also shown that matching prior knowledge, or expectation can affect neural responses to speech and representations of acoustic and phonetic features. However, these findings were observed under a fixed degradation level, close to our lowest-clarity, 3-channel condition (see also Di Liberto et al., 2018a). Thus, it is unclear whether the enhanced neural representations observed in these earlier studies reflect computations of sharpened signals or prediction errors – both types of representational scheme can accommodate the finding that prior knowledge enhances neural representations for low clarity speech (see Figure 1D). By manipulating degradation level alongside prior knowledge, we could test for the presence of interactive effects that distinguish neural representations of prediction errors from sharpened signals. We observed that matching prior knowledge enhances neural representations of low-clarity speech but leads to suppression of neural representations for high-clarity speech. This interaction is uniquely consistent with prediction error representations (Blank and Davis, 2016).

Other work has taken a different approach to investigating how prior knowledge affects speech processing. These previous studies capitalized on the fact that in natural speech, certain speech sounds are more or less predictable based on the preceding sequence of phonemes (e.g. upon hearing “capt-”, “-ive” is more predictable than “-ain” because “captain” is a more frequent English word than “captive”). Using this approach, MEG studies demonstrate correlations between the magnitude of neural responses and the magnitude of phoneme prediction error (i.e. phonemic surprisal, Brodbeck et al., 2018; Donhauser and Baillet, 2020). However, this previous work did not test for sharpened representations of speech; leaving open the question of whether expectations are combined with sensory input by computing prediction errors or sharpened signals. Indeed, this question may be hard to address using natural listening paradigms since many measures of predictability are highly correlated (e.g. phonemic surprisal and lexical uncertainty, Brodbeck et al., 2018; Donhauser and Baillet, 2020). Here, by experimentally manipulating listeners’ predictions and the quality of sensory input, we could test for the statistical interaction between prior knowledge and sensory detail that most clearly distinguishes prediction errors from sharpened representations. Future work in which this approach is adapted to continuous speech could help to establish the role of predictive computations in supporting natural speech perception and comprehension.

## Methods

### Participants

Twenty-one (12 female, 9 male) right-handed participants were tested after being informed of the study’s procedure, which was approved by the Cambridge Psychology Research Ethics Committee. All were native speakers of English, aged between 18 and 40 years (mean = 22, SD = 2) and had no history of hearing impairment or neurological disease based on self-report.

### Spoken stimuli

468 monosyllabic words were presented to each participant in spoken or written format drawn randomly from a larger set of monosyllabic spoken words. The spoken words were 16-bit, 44.1 kHz recordings of a male speaker of southern British English and their duration ranged from 372 to 903 ms (mean = 591, SD = 78). The amount of sensory detail in speech was varied using a noise-vocoding procedure (Shannon et al., 1995), which superimposes the temporal-envelope from separate frequency regions in the speech signal onto white noise filtered into corresponding frequency regions. This allows parametric variation of spectral detail, with increasing numbers of channels associated with increasing intelligibility. Vocoding was performed using a custom Matlab script (The MathWorks Inc.), using 3, 6, or 12 spectral channels spaced between 70 and 5000 Hz according to Greenwood’s function (Greenwood, 1990). Envelope signals in each channel were extracted using half-wave rectification and smoothing with a second-order low-pass filter with a cut-off frequency of 30 Hz. The overall RMS amplitude was adjusted to be the same across all audio files.

Each spoken word was presented only once in the experiment so that unique words were heard on all trials. The particular words assigned to each condition were randomized across participants. Before the experiment, participants completed brief practice sessions each lasting approximately five minutes that contained all conditions but with a different corpus of words to those used in the experiment. Stimulus delivery was controlled with E-Prime 2.0 software (Psychology Software Tools, Inc.).

### Procedure

Participants completed a modified version of the clarity rating task previously used in behavioural and MEG studies combined with a manipulation of prior knowledge (Davis et al., 2005; Hervais-Adelman et al., 2008; Sohoglu et al., 2012). Speech was presented with 3, 6 or 12 channels of sensory detail while prior knowledge of speech content was manipulated by presenting mismatching or matching text before speech onset (see Figure 1A). Written text was composed of black lowercase characters presented for 200 ms on a gray background. Mismatching text was obtained by permuting the word list for the spoken words. As a result, each written word in the Mismatching condition was also presented as a spoken word in a previous or subsequent trial and vice versa.

Trials commenced with the presentation of a written word, followed 1050 (± 0-50) ms later by the presentation of a spoken word (see Figure 1A). Participants were cued to respond by rating the clarity of each spoken word on a scale from 1 (‘Not clear’) to 4 (‘Very clear’) 1050 (±0-50) ms after speech onset. The response cue consisted of a visual display of the rating scale and responses were recorded using a four-button box from the participant’s right hand. Subsequent trials began 850 (±0-ms after participants responded.

Manipulations of sensory detail (3/6/12 channel speech) and prior knowledge of speech content (Mismatching/Matching) were fully crossed, resulting in a 3 × 2 factorial design with 78 trials in each condition. Trials were randomly ordered during each of three presentation blocks of 156 trials.

### Data acquisition and pre-processing

Magnetic fields were recorded with a VectorView system (Elekta Neuromag, Helsinki, Finland) containing two orthogonal planar gradiometers at each of 102 positions within a hemispheric array. Data were also acquired by magnetometer and EEG sensors. However, only data from the planar gradiometers were analyzed as these sensors have maximum sensitivity to cortical sources directly under them and therefore are less sensitive to noise artifacts (Hämäläinen, 1995).

MEG data were processed using the temporal extension of Signal Source Separation (Taulu et al., 2005) in Maxfilter software to suppress noise sources, compensate for motion, and reconstruct any bad sensors. Subsequent processing was done in SPM8 (Wellcome Trust Centre for Neuroimaging, London, UK), FieldTrip (Donders Institute for Brain, Cognition and Behaviour, Radboud University Nijmegen, the Netherlands) and NoiseTools software (http://audition.ens.fr/adc/NoiseTools) implemented in Matlab. The data were subsequently highpass filtered above 1 Hz and downsampled to 80 Hz (with anti-alias lowpass filtering).

### Linear regression

We used ridge regression to model the relationship between speech features and MEG responses (Pasley et al., 2012; Ding and Simon, 2013; Di Liberto et al., 2015; Holdgraf et al., 2017; Brodbeck et al., 2018). Prior to model fitting, the MEG data were epoched at matching times as the feature spaces (see below), outlier trials removed and the MEG time-series *z*-scored such that each time-series on every trial had a mean of zero and standard deviation of 1. We then used the mTRF toolbox (Crosse et al., 2016; https://sourceforge.net/projects/aespa) to fit two types of linear model.

The first linear model was an encoding model that mapped from the stimulus features to the neural time-series observed at each MEG sensor and at multiple time lags:

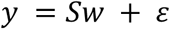

where *y* is the neural time-series recorded at each planar gradiometer sensor, *S* is an N_samples_ by N_features_ matrix defining the stimulus feature space (concatenated over different lags), w is a vector of model weights and *ε* is the model error at each timepoint. Model weights for positive and negative lags capture the relationship between the speech feature and the neural response at later and earlier timepoints, respectively. Encoding models were fitted with lags from -100 to 300 ms but for model prediction purposes (see below), we used a narrower range of lags (0 to 250 ms) to avoid possible artefacts at the earliest and latest lags (Crosse et al., 2016).

Our encoding models tested four feature spaces (shown in Figure 2), derived from the original clear versions of the spoken stimuli (i.e. prior to noise-vocoding):

1. Broadband envelope, obtained by summing the envelopes across the spectral channels in a 24-channel noise-vocoder. The center frequencies of these 24 spectral channels were Greenwood spaced: 86, 120, 159, 204, 255, 312, 378, 453, 537, 634, 744, 869, 1011, 1172, 1356, 1565, 1802, 2072, 2380, 2729, 3127, 3579, 4093 and 4678 Hz (see Greenwood, 1990). Envelopes were extracted using half-wave rectification and low-pass filtering at 30 Hz.
2. Spectrogram, which was identical to the Envelope feature space but with the 24 individual spectral channels retained.
3. Spectrotemporal modulations captured regular fluctuations in energy across the frequency and time axes of the spectrogram and was obtained using the NSL toolbox in Matlab (http://nsl.isr.umd.edu/downloads.html). This toolbox first computes a 128-channel ‘auditory’ spectrogram to simulate processing in the cochlea and midbrain (with constant Q and logarithmic center frequencies between 180 and 7040 Hz), before filtering with 2D wavelet filters tuned to spectral modulations of 0.5, 1, 2, 4 and 8 cycles per octave and temporal modulations of 1, 2, 4, 8 and 16 Hz. All other parameters were set to the default (frame length = 8 ms, time constant = 8 ms and no nonlinear compression). Note that temporal modulations can be positive or negative, reflecting the direction of frequency sweeps (i.e. upward versus downward; Elliott and Theunissen, 2009). This default representation is a very high-dimensional feature space: frequency x spectral modulation x temporal modulation x temporal modulation direction with 128 × 5 x 5 × 2 = 6400 dimensions, each represented for every 8 ms time sample in the speech file. We therefore averaged over the 128 frequency channels and positive-and negative-going temporal modulation directions. The resulting feature space was a time-varying representation of spectrotemporal modulation content comprised of 25 features (5 spectral modulations x 5 temporal modulations).
4. Phonetic features. The final feature space comprised 13 phonetic features describing the time-varying phonetic properties of speech including vowel backness, voicing, place and manner of articulation. We first used a forced-alignment algorithm included as part of BAS software (Kisler et al., 2017; https://clarin.phonetik.uni-muenchen.de/BASWebServices/interface/WebMAUSGeneral) to estimate the onset time of each phoneme. We then converted phoneme representations into the following articulatory phonetic features: Voiced, Unvoiced, Bilabial, Labiodental/Dental, Alveolar, Velar, Plosive, Nasal, Fricative, Liquid, Front, Central and Back (International Phonetic Association, 1999). Between the onset time of each phoneme and the onset time of the subsequent phoneme, the corresponding phonetic features were set to 1 and non-corresponding phonetic features to 0. Phonetic features that occurred infrequently in our stimulus set were excluded or merged together (labiodental and dental features). For dipthong vowels and affricate consonants, we averaged the feature vectors for the component speech sounds.

In addition to the four feature spaces above, we also tested more complex combinations of feature spaces that previous work has shown to be good predictors of neural responses. The first of these combined the spectrogram feature space above with the phonetic feature space (Di Liberto et al., 2015). The second combined the spectrogram with the half-wave rectified derivative of the spectrogram to capture spectral onsets (Daube et al., 2019).

All feature spaces were downsampled to 80 Hz to match the MEG sampling rate and the acoustic feature spaces (i.e. 1, 2, 3 above) were *z*-score transformed such that each time-series on every trial had a mean of zero and standard deviation of 1. For each trial/word, all feature spaces were zero-padded before speech onset to match the duration of the longest negative lag used for model fitting (i.e. 100 ms). Similarly, zero-padding was added after speech offset to match the duration of the longest positive lag (i.e. 300 ms). This enabled information at the beginning and end of each spoken word to inform the model fits for negative and positive lags, respectively.

The second linear model was a decoding model that mapped in the opposite direction from the neural time-series back to speech spectrotemporal modulations (since analysis of encoding models showed this feature space to be most predictive of neural responses). Prior to model fitting, to remove redundant dimensions and reduce computation time, the data from the 204 planar gradiometers were transformed into principal components and the first 50 components retained. The model was therefore as follows:

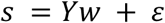

Here *s* is a vector expressing the time-varying spectrotemporal modulations (one vector for each of the 25 spectrotemporal modulations), *Y* is an N_samples_ by N_components_ matrix of MEG data (concatenated over different lags), *w* is a vector of model weights and *ε* is the model error at each timepoint. Decoding models were fitted with lags from -300 to 100 ms and from -250 to 0 ms for model prediction purposes. Here negative and positive lags index the relationship between the neural response and speech at earlier and later timepoints, respectively.

To control for overfitting, we used ridge regression and varied the lambda parameter (over 17 values as follows: 2^0^, 2^1^, 2^3^, … 2^20^) which varies the degree to which strongly positive or negative weights are penalized (Crosse et al., 2016). This lambda parameter was optimized using a leave-one-trial-out cross-validation procedure. For each feature space, sensor, condition and participant, we fitted the models using data from all but one trial. We then averaged the model weights across trials and used the result to predict the MEG response (for the encoding models) or speech spectrotemporal modulations (for the decoding model) of the left-out trial. We computed model accuracy as the Pearson correlation between the predicted and observed data and repeated this procedure such that model accuracies were obtained for all trials. Before optimizing lambda, model accuracies were averaged across trials, and also across sensors (encoding models) or modulations (decoding model). The optimal lambda value was then selected as the mode of the model accuracy distribution over participants and conditions (Holdgraf et al., 2016). When testing the complex models that combined feature spaces, lambda was optimized for each component feature space independently resulting in a search space of 17 × 17 = 289 lambda values (i.e. “banded” ridge regression; Nunez-Elizalde et al., 2019).

### Statistical analysis

In addition to using model accuracies to control for overfitting (see ‘Model fitting’ section above), we also compared model accuracies for different feature spaces (to determine which feature space was best represented in MEG responses) and for different conditions (to examine how feature representations were modulated by our experimental manipulations). In the case of the encoding models, we averaged model accuracies over trials and over the 20 sensors with the highest model accuracies (in each hemisphere separately). For the decoding model, model accuracies were averaged over trials. The resulting data were then entered into repeated measures ANOVAs or *t*-tests to examine differences in model accuracy between feature spaces or conditions.

To visualize the weights of the spectrotemporal modulation encoding model, or ‘temporal response functions (TRFs)’, we used Denoising Source Separation (DSS) to transform the TRFs into a set of linear components (spatial filters) ordered by their consistency over trials (de Cheveigné and Parra, 2014). The first 3 DSS components (i.e. the 3 most consistent components) were retained and projected back into sensor-space. When applying DSS, the covariance matrices were computed by summing the covariances over features and conditions (for each feature space separately). Following DSS, we averaged the TRFs across trials and computed the Root Mean Square (RMS) amplitude across all left hemisphere sensors.

## Acknowledgements

This research was supported by the Medical Research Council (MC-A060-5PQ80 to M.H.D.). We are grateful to Maarten van Casteren, Clare Cook and Lucy MacGregor for their excellent technical support during data collection.

